# Genome-wide prediction of disease variants with a deep protein language model

**DOI:** 10.1101/2022.08.25.505311

**Authors:** Nadav Brandes, Grant Goldman, Charlotte H. Wang, Chun Jimmie Ye, Vasilis Ntranos

**Affiliations:** Division of Rheumatology, Department of Medicine, UCSF, University of California, San Francisco; Department of Epidemiology and Biostatistics, Institute of Computational Health Sciences, University of California, San Francisco; Diabetes Center, University of California, San Francisco; Biomedical Sciences Graduate Program, University of California, San Francisco; Parker Institute for Cancer Immunotherapy, Chan Zuckerberg Biohub, San Francisco

## Abstract

Distinguishing between damaging and neutral missense variants is an ongoing challenge in human genetics, with profound implications for clinical diagnosis, genetic studies and protein engineering. Recently, deep-learning models have achieved state-of-the-art performance in classifying variants as pathogenic or benign. However, these models are currently unable to provide predictions over all missense variants, either because of dependency on close protein homologs or due to software limitations. Here we leveraged ESM1b, a 650M-parameter protein language model, to predict the functional impact of human coding variation at scale. To overcome existing technical limitations, we developed a modified ESM1b workflow and functionalized, for the first time, all proteins in the human genome, resulting in predictions for all ∼450M possible missense variant effects. ESM1b was able to distinguish between pathogenic and benign variants across ∼150K variants annotated in ClinVar and HGMD, outperforming existing state-of-the-art methods. ESM1b also exceeded the state of the art at predicting the experimental results of deep mutational scans. We further annotated ∼2M variants across ∼9K alternatively-spliced genes as damaging in certain protein isoforms while neutral in others, demonstrating the importance of considering all isoforms when functionalizing variant effects. The complete catalog of variant effect predictions is available at: https://huggingface.co/spaces/ntranoslab/esm_variants.

## Introduction

Determining the phenotypic consequences of genetic variants is a key challenge in human genetics [1–4]. While most genetic variants associated with common human disease occur in non-coding regions of the genome, variants that affect the amino-acid (AA) sequence of proteins are enriched amongst variants associated with common human disease and cause many rare Mendelian and somatic disorders [5–8]. Coding variants are also of special interest because the mechanisms by which they act are better understood, and they are more therapeutically actionable. The vast majority of coding variants are missense variants that substitute one AA with another along the protein sequence [9]. However, distinguishing between damaging variants, which disrupt the biological function of the protein, from neutral variants, which have no functional effects, remains challenging. Furthermore, the same genetic variant can be present in multiple isoforms but only have a damaging effect in a subset of them, depending on interactions with other parts of the protein sequence. As a result, most missense variants are still labeled as Variants of Uncertain Significance (VUS), preventing many patients from receiving a clear medical diagnosis [2, 10].

Variant functionalization can be performed either experimentally or computationally. Experimental approaches such as deep mutational scans [11] and Perturb-Seq [12] can measure cellular phenotypes (e.g., gene expression or reporter activity) across hundreds to thousands of variants simultaneously. However, their current limited scale precludes genomewide experiments, and the specific cellular phenotypes measured are imperfect proxies for the clinical phenotype of interest (e.g., disease risk or therapeutic response) [13, 14]. In contrast, computational methods that learn the biophysical properties or evolutionary constraints of protein sequences are theoretically scalable to all coding variants [15–17]. Most existing computational algorithms for variant functionalization are supervised, trained on databases of pathogenic versus benign variants [10]. Recently, an unsupervised deep-learning method named EVE, which predicts variant effects directly from amino-acid sequences without training on labeled data, has shown to outperform supervised methods [4]. EVE builds a model based on a multiple sequence alignment (MSA) separately for each human protein. EVE currently provides predictions for only ∼15% of human proteins, and because of its reliance on MSA coverage, can only predict the effects of well-aligned residues in each protein. Moreover, since alternative isoforms of the same human protein share the same homologs, it is not clear whether EVE and other homology-based methods are capable of distinguishing between variant effects on different isoforms.

A different deep-learning approach to functionalizing variants uses protein language models, a technique originally developed for natural language processing, which has been successfully applied to many protein prediction tasks [18–23]. Protein language models are trained on a large set of protein sequences that have been selected throughout evolution (e.g. from UniProt [24]), thereby learning which sequence variations are more or less likely to occur. Importantly, protein language models don’t require explicit homology information, they are not gene-specific, and the same model applies for any possible AA sequence. It has been shown that protein language models implicitly learn to recognize many aspects of protein structure and function, such as secondary structure, long-distance interactions (e.g. disulfide bonds), post-translational modifications and binding sites. ESM1b [19], a 650M-parameter model trained on ∼250M protein sequences (**Fig. 1A**), was demonstrated to predict, without further training, variant effects correlated with the results of mutational scanning experiments [25].

**Figure 1.**
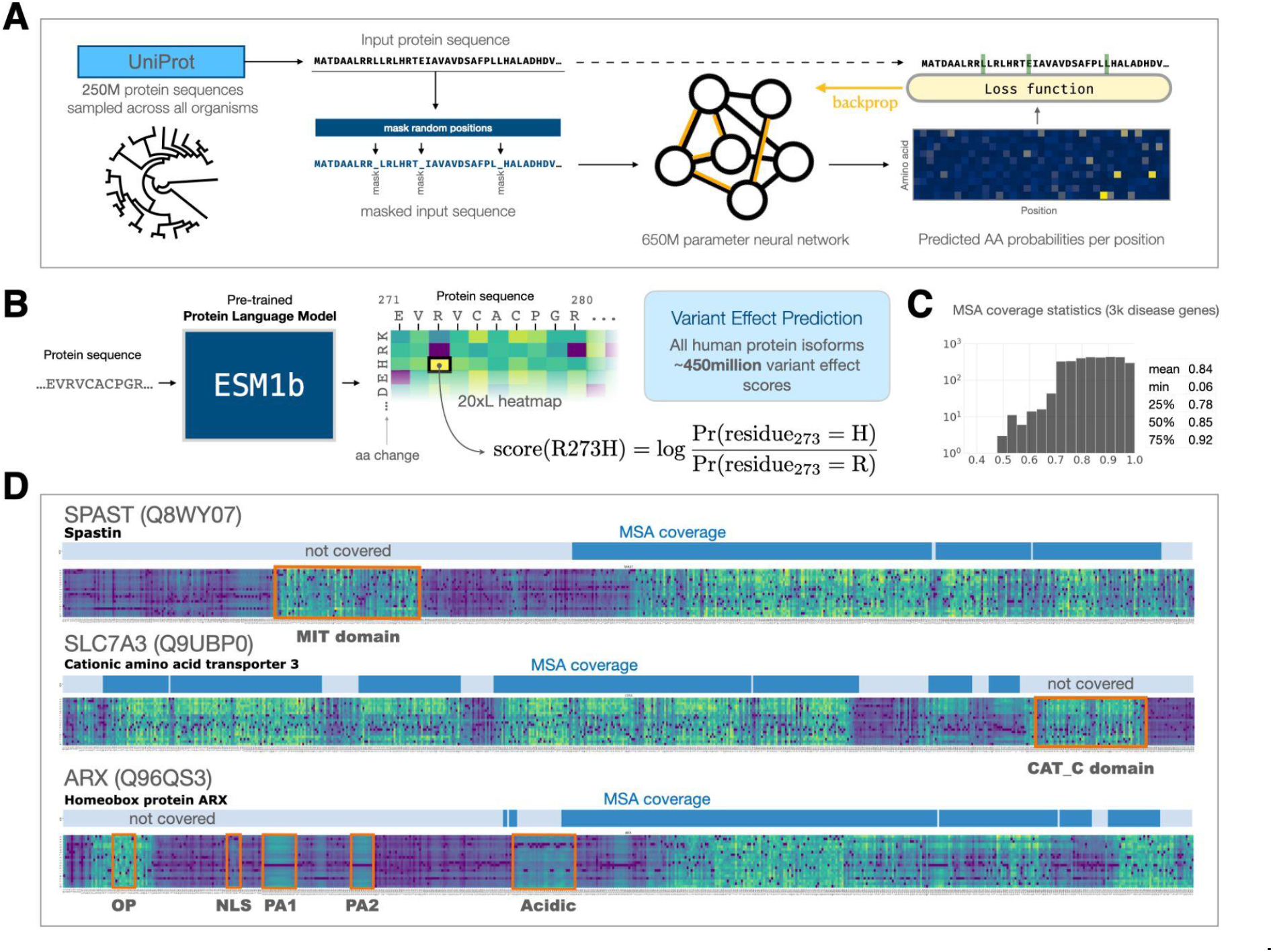
ESM1b predicts variant effects without homology coverage. (**A**) ESM1b is a 650M-parameter protein language model trained on 250M protein sequences across all organisms. The model was trained via the masked language modeling task, where random residues are masked from input sequences and the model has to predict the correct amino acid at each position (including the missing residues). (**B**) Illustration of the ESM1b model’s input (an amino-acid sequence) and output (log likelihood ratio of effect scores for all possible missense variants). (**C**) The distribution of MSA coverage (i.e. the fraction of a protein’s residues that are aligned) across ∼3K disease-related proteins. (**D**) Examples of the model’s capacity to detect protein domains and functional regions across three human proteins: SPAST, SLC7A3 and ARX. Each heatmap visualizes the log-likelihood ratio scores across all *20xL* possible missense variants (where *L* is the protein length). Protein domains without MSA coverage are highlighted in red.

Despite these advantages, there are several technical limitations of ESM1b that limit its application for variant functionalization. First, ESM1b is technically limited to sequence inputs of up to 1,022aa in length, which is inadequate for handling the full-length sequences of about 12% of the human protein isoforms. Second, while ESM1b variant effect predictions have been evaluated on cellular phenotypes across 32 genes (10 from human) [25], the performance of the method in predicting the clinical impact of human coding variants across the genome is unknown. Finally, ESM1b is implemented in a computational framework that requires software engineering proficiency, deep-learning expertise, and specialized hardware (high-memory GPUs), which together create a technical barrier for its broad adoption by geneticists and clinicians.

Here, we sought to lift these barriers and enable ESM1b to inform clinical diagnosis and human genetics research at a genome-wide scale. To this end, we implemented a workflow generalizing ESM1b to protein sequences of any length, and used it to predict the effects of all ∼450M possible single AA substitutions across all 42,336 protein isoforms in the human genome. Our ESM1b-based workflow outperforms EVE as a classifier of variant pathogenicity (as annotated by ClinVar [10] and HGMD [26]) and as a predictor of deep mutational scanning experiments. We further demonstrate the capacity of ESM1b to assess variant effects in the context of different protein isoforms. Using our workflow, we identified variants predicted to be isoform sensitive in 85% of alternatively spliced genes. These isoform-sensitive variants, which are not captured by traditional approaches based solely on the inclusion or exclusion of entire exons, are enriched near splice sites and in genes containing known alternatively spliced domains. Finally, we created a web portal that allows researchers to query, visualize and download variant effect predictions for all protein isoforms in the human genome. This complete catalog is accessible at: https://huggingface.co/spaces/ntranoslab/esm_variants.

## Results

We developed and applied a modified ESM1b workflow on all 42,336 known human protein isoforms, obtaining a complete catalog of all ∼450M possible missense variant effects in the human genome. The effect score of each missense variant (in the context of a protein sequence) is defined as the log likelihood ratio (LLR) between the variant and the wild-type amino acid (**Fig. 1B**). Compared to homology-based models that are currently available for a small subset of human proteins and cover only specific residues in those proteins (e.g. 84% of the residues in 15% of human proteins in the case of EVE; **Fig. 1C**), ESM1b produces variant effect scores for every possible missense variant in every protein.

Protein regions that are more functionally sensitive according to ESM1b (i.e. with many variants predicted to be damaging) often align with known protein domains and harbor disease-associated variants (**Fig. 1D**). For example, as illustrated for *SPAST, SLC7A3,* and *ARX*, such protein domains sometimes reside outside of MSA coverage, making predictions of variants in these domains unavailable to homology-based models (**Fig. 1D**). For these genes, we also observed variants associated with human disease residing in regions not covered by MSA. For example, the MIT domain in *SPAST* contains 4 missense variants (E112K, R115C, N184T, and L195V) that have been observed in patients with hereditary spastic paraplegias, and further supported as causal by experimental evidence [27]. Likewise, the CAT C domain in *SLC7A3* contains a missense variant (S589T) identified in males with autism spectrum disorder and intellectual disability [28]. Similarly, multiple domains in *ARX* outside of MSA coverage (highlighted in **Fig. 1D**) contain genetic variation implicated with intellectual disability [29–32].

To evaluate how well ESM1b predicts the clinical consequences of variants, we compared the effect scores predicted by the model between pathogenic and benign variants in two datasets. The first dataset contains variants annotated as pathogenic or benign according to ClinVar [10]. The second dataset contains variants annotated as disease causing in HGMD [26] and benign variants defined by allele frequency greater than 1% in gnomAD [9]. Indeed, the distribution of pathogenic variant effect scores is dramatically different from the distribution for benign variants in both datasets. Moreover, we observe consistent effect scores for both variant labels (pathogenic and benign) between the two datasets (ClinVar and HGMD/gnomAD; **Fig. 2A**), suggesting that the predictions are well calibrated across genes. Using an LLR threshold of -7.5 (near the intersection of the two distributions) to distinguish between pathogenic and benign variants yields a true positive rate of 81% and true negative rate of 82% across both datasets.

**Figure 2.**
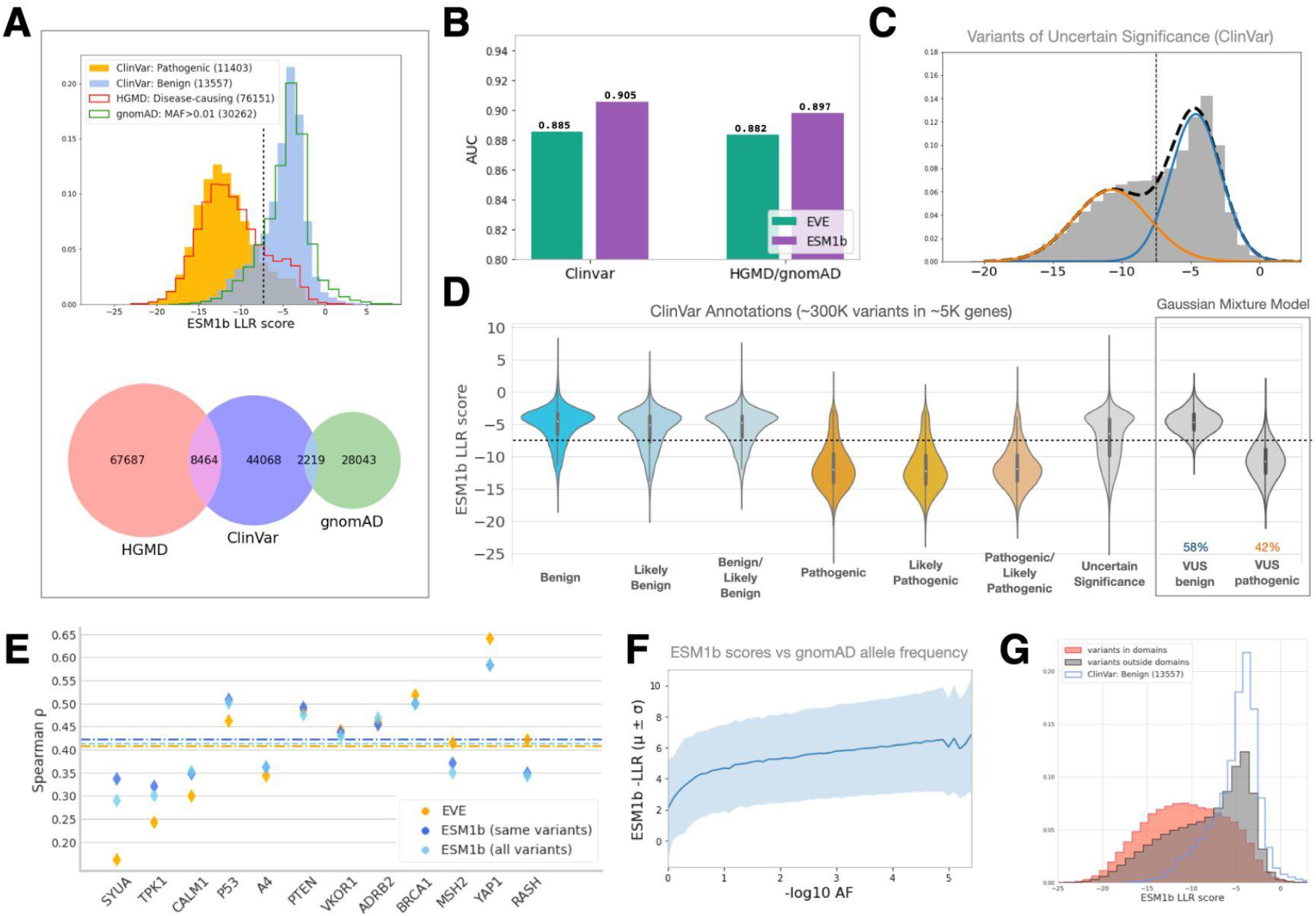
ESM1b is suitable for genome-wide disease prediction of coding variants. (**A**) Top panel: the distribution of ESM1b scores across two sets of variants that are assumed to be mostly pathogenic (“ClinVar: Pathogenic” and “HGMD: Disease-causing”) and two sets of variants assumed to be mostly benign (“ClinVar: Benign:” and “gnomad: MAF>0.01”). Bottom panel: venn diagram of the variants extracted from HGMD, ClinVar and gnomAD. (**B**) Comparison between ESM1b and EVE in their capacity to distinguish between pathogenic and benign variants (measured by global ROC-AUC scores), as labeled by ClinVar (36,537 variants in 2,765 unique genes) or HGMD/gnomAD (30,497 variants in 1,991 unique genes). (**C**) The distribution of ESM1b scores across ClinVar variants of uncertain significance (VUS), decomposed as a mixture of two Gaussian distributions capturing variants predicted as more likely pathogenic (orange) or more likely benign (blue). (**D**) The distribution of ESM1b scores across all common ClinVar labels, including the two Gaussian components from (C). (**E**) Comparison between ESM1b and EVE on deep mutational scan datasets covering 12 human genes (**Supplementary Table S1**). Performance is measured by the Spearman’s correlation between each model’s scores and the experimental scores. Average performance of each of the models is marked by a dashed line. Since ESM1b can process all missense variants (unlike EVE which only assigns scores for a subset of them), the performance of ESM1b is shown either for all variants (“all variants”) or the subset of variants with EVE scores (“same variants”). (**F**) Average ESM1b score (and standard-deviation) as a function of allele frequency over all gnomAD missense variants. (**G**) The distribution of ESM1b scores across variants in annotated protein domains (red) vs. variants outside domains (gray). The distribution of benign variants (as in (A)) is shown for reference.

Next, we compared the performance of ESM1b and EVE as classifiers of variant pathogenicity. On ClinVar, ESM1b obtains a ROC-AUC score of 0.905 for distinguishing between the 19,925 pathogenic and 16,612 benign variants (across 2,765 unique genes), compared to 0.885 for EVE. Similarly, ESM1b obtains a ROC-AUC score of 0.897 at discerning between the 27,754 variants labeled by HGMD as disease causing and the 2,743 common missense variants in gnomAD (across 1,991 unique genes), compared to 0.882 for EVE (**Fig. 2B**). Controlling for the false positive rate using the standard 0.05 threshold, ESM1b obtains a true positive rate of 60% compared to 49% for EVE over ClinVar, and 61% compared to 51% over HGMD/gnomAD (**Supplementary Fig. S1**).

Having established the high accuracy of ESM1b as a classifier of variant pathogenicity, we sought to predict the effects of VUS currently annotated in ClinVar. To that end, we modeled the distribution of ESM1b scores across VUS as a Gaussian mixture with two components (**Fig. 2C**). Indeed, these two fitted distributions align well with the distributions for annotated pathogenic and benign variants (**Fig. 2D**). According to this model, we estimate that about 58% of VUS in ClinVar are benign and about 42% are pathogenic.

Next, we compared ESM1b and EVE on experimental measurements of deep mutational scans conducted across 21 assays in 12 human genes (**Supplementary Table S1**). On average, ESM1b obtains a Spearman’s correlation of 0.422 between its effect scores and the experimental scores, compared to 0.407 for EVE (**Fig. 2E**).

We further assessed the functional consequences of ESM1b predictions in two analyses. First, we found that ESM1b scores track well with allele frequency; the more common a variant is, the less likely it is to obtain ESM1b scores considered damaging (**Fig. 2F**). Second, as illustrated by individual examples (**Fig. 1D**), the distribution of ESM1b scores for variants residing within domains is more damaging, while the distribution for variants residing outside of domains is more similar to benign variants (**Fig. 2G**).

As a protein language model, ESM1b evaluates each variant in the context of the entire protein sequence provided as input. This provides an opportunity to assess how the contextual sequence of alternative isoforms influences the model’s effect predictions. It is possible that the same variant could be damaging in the context of some protein isoforms but not others, for example due to interaction with alternatively spliced domains (**Fig. 3A**). For example, P53 has a shorter isoform with both the N- and C-termini truncated. By generating variant effect predictions for the primary isoform and this alternative isoform independently, we found that 170 variants (mostly near the splice junctions) obtain substantially different scores (LLR diff > 4) between the two isoforms, including three ClinVar variants annotated as VUS (**Fig. 3B**).

**Figure 3.**
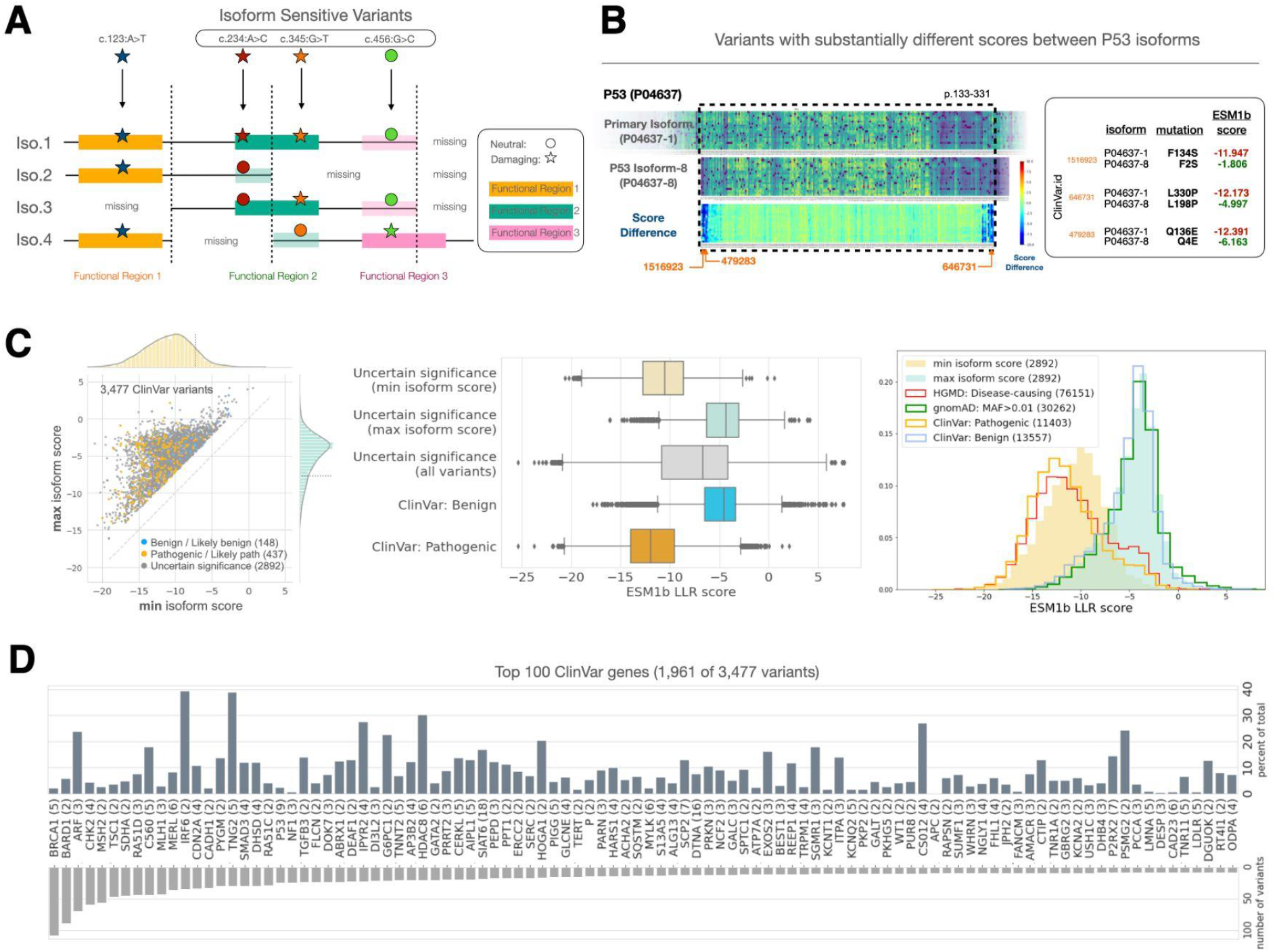
ESM1b predictions in clinically relevant genes depend on the isoform context. (**A**) The consequences of variants (e.g. damaging vs. neutral) can depend on the isoform context. (**B**) Comparison of the primary and one of the alternative isoforms of P53. Three specific variants are detailed. (**C**) Left panel: all 3,477 ClinVar variants with highly variable ESM1b effect scores (defined by *std>2*) across different isoforms. Center panel: the lowest and highest isoform scores predicted for VUS from the left panel (top 2 boxes), compared to the mean scores (across isoforms) of VUS, benign or pathogenic variants (as in Fig. 2D; bottom 3 boxes). Right panel: The distribution of the lowest and highest isoform scores predicted for VUS from the left panel, compared to the distributions for pathogenic or benign variants from ClinVar, HGMD and gnomAD (as in Fig. 2A). Across the entire panel, the number of variants associated with each category is shown in parentheses. (**D**) The top 100 ClinVar genes with the largest number of variants with highly variable effect scores (as in (C)). In parentheses: the number of annotated isoforms of each gene.

We next searched for missense variants over the ClinVar dataset with substantially different predicted effects between isoforms. Specifically, we found 3,477 variants with LLR standard deviation greater than 2 across all isoforms (**Fig. 3C**). Of these 3,477 variants, 148 (4%) are benign or likely benign, 437 (13%) are pathogenic or likely pathogenic, and 2,892 (83%) are annotated as VUS. Interestingly, considering these 2,892 VUS in the context of the isoform predicted to be most damaging provides an effect score distribution similar to that of pathogenic variants, while considering the isoform predicted to be least damaging provides a distribution similar to benign variants (**Fig. 3C**). Like P53, many clinically important human genes have a large number of ClinVar variants with high variance in ESM1b scores across isoforms, including *BRCA1, IRF6* and *TGFB3* (**Fig. 3D**).

Beyond the ∼5k ClinVar genes, we further examined isoform-specific effects genome-wide across all possible missense variants and 20,360 coding human genes. To this end, we define a variant to be isoform-sensitive if i) it is predicted by ESM1b as likely bening (LLR>-7) in at least one isoform, ii) it is predicted as likely pathogenic (LLR<-8) in at least one isoform, and iii) these two predictions are substantially different (LLR difference > 4). We found ∼1.8M such variants across ∼9K genes, which is 85% of all genes with annotated alternative isoforms (**Fig. 4A**). Isoform sensitive variants are more likely to occur near splice junctions and in genes with protein domains disrupted by splicing events, as opposed to domains that are either included intact or removed entirely across alternative isoforms (**Fig. 4B**).

**Figure 4.**
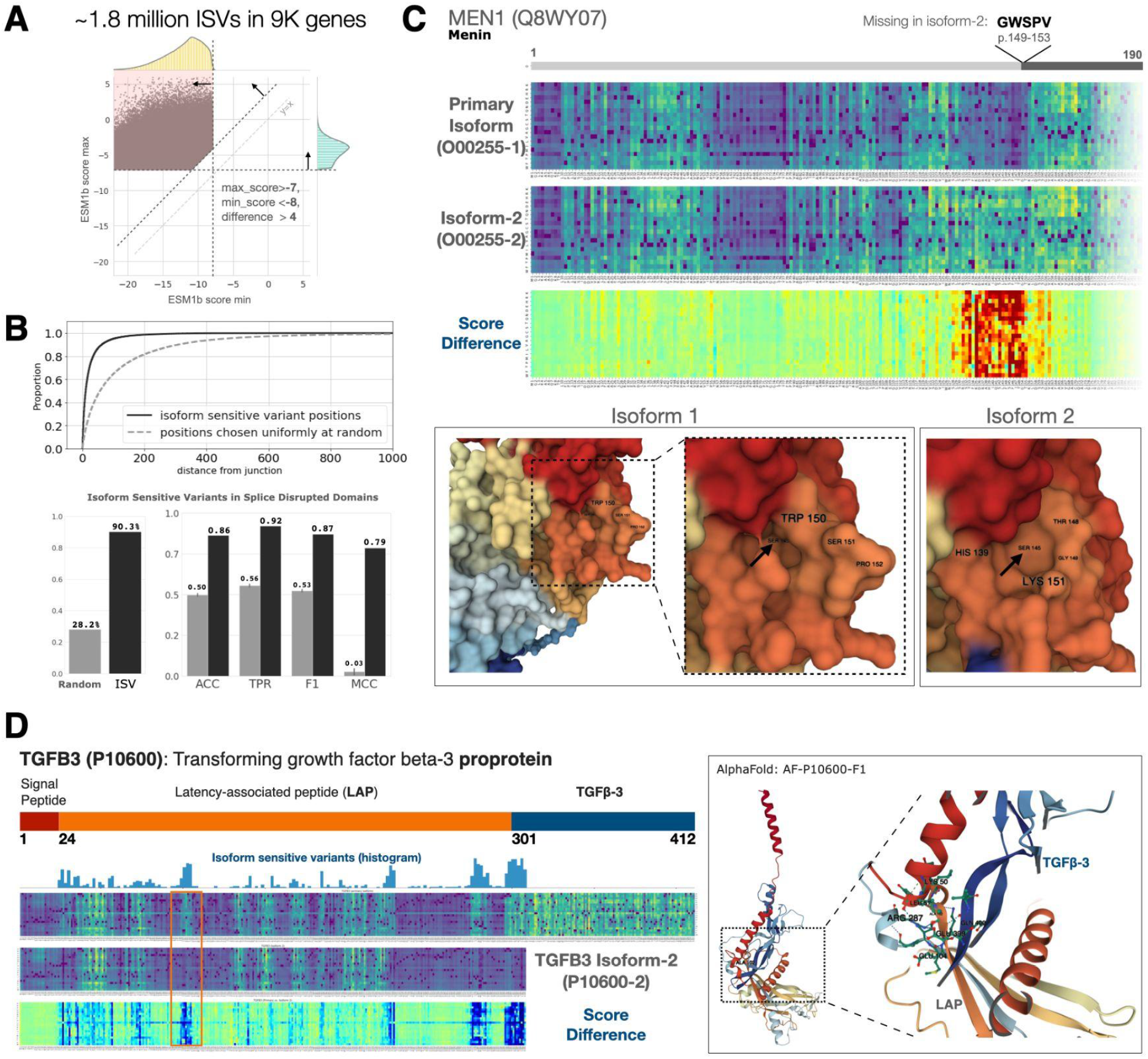
ESM1b can detect isoform-specific variant effects. (**A**) ∼1.8M missense variants across ∼9K genes in the human genome are “isoform sensitive”, defined by: i) maximum ESM1b score (across isoforms) >-7, ii) minimum score <-8, iii) difference between minimum and maximum score >4. (**B**) Top panel: isoform sensitive variants are closer to splice junction than would be expected at random. Bottom-left panel: isoform sensitive variants in genes with domains containing splice junctions: 90.31% vs. 28.21% expected at random. Bottom-right panel: metrics of predicting whether genes contain domains disrupted by splice junction given whether or not they contain isoform sensitive variants. (**C**) An example of a small splicing effect (excision of 5 amino acids from the primary isoform of the *MEN1* protein) leading to dramatic changes in the predicted effects of variants in a much larger region. Bottom panel: AlphaFold structural predictions of the two isoforms. (**D**) An example of alternative splicing leading to a distant effect in the *TGFB3* proprotein. Exclusion of the TGFβ-3 chain in an alternative isoform of the proprotein leads to a region at the beginning of the LAP chain (marked by orange) to lose its sensitivity to missense variants. Right panel: AlphaFold prediction of the binding of the two chains showing these two regions to be close to one another in 3D structure.

We found evidence of splicing events that dramatically influence the predicted variant effects. For example, the second isoform of MEN1, a tumor suppressor gene involved in many types of cancer, differs from the primary isoform by the deletion of only five amino acids at positions 149-153 due to an alternative donor site at exon 1. Based on ESM1b scores, we predict that this deletion introduces a 30aa-long functional region preceding that location, making variants in this region more prone to be considered damaging (**Fig. 4C**). Indeed, multiple studies have reported missense variants in that region to be associated with cancer, including: H139D [33], H139P [34], H139R [35], H139Y [36], F144V [37] and I147F [38]. A study from 2017 found that the second MEN1 isoform displays aberrant expression in human hepatocellular carcinoma and cholangiocarcinoma relative to matched normal tissue in TCGA; however, the authors remarked that functional differences between the two isoforms were yet to be characterized [39]. The functional role of the 5 aa deletion is further supported by the predicted 3D structures of the two protein isoforms [40], which differ by a small pocket on the protein’s surface (**Fig. 4C**).

Another example of isoform-sensitive variants can be seen in the TGFB3 (transforming growth factor beta-3) proprotein. This proprotein is cleaved into two separate chains, LAP and TGFβ-3, which together form a dimer. However, an alternative isoform of TGFB3 is lacking the TGFβ-3 chain, causing many missense variants in the LAP chain to no longer be deemed damaging by ESM1b. The region with isoform-sensitive variants in the LAP chain is more than 200 residues away from the alternatively spliced TGFβ-3 chain. However, according to AlphaFold [40], they are in close contact in 3D space (**Fig. 4D**).

## Discussion

Using a state-of-the-art protein language model ESM1b, we have compiled a complete catalog of predicted effect scores for all possible missense variants in the human genome (https://huggingface.co/spaces/ntranoslab/esm_variants). A comprehensive evaluation of the effect scores demonstrates that ESM1b predictions exceed the performance of EVE at distinguishing between pathogenic and benign variants across ClinVar and HGMD/gnomAD and in predicting the effects reported by deep mutational scan assays. Previous analysis has established EVE as state of the art for variant functionalization based on these two benchmarks, outperforming 7 unsupervised and 8 supervised variant effect prediction methods [4], including CADD [17], SIFT [15] and Polyphen2 [16]. We have also demonstrated that ESM1b can recognize protein domains and other functional regions, including those outside MSA coverage (**Fig. 1D**), and that its predicted effect scores are correlated with allele frequency (**Fig. 2F**).

As an unsupervised model that doesn’t rely on an explicit model of homology, ESM1b avoids many of the potential pitfalls of existing methods. Compared to supervised methods, when validating it on clinical (e.g. ClinVar and HGMD) or population-genetics (e.g. gnomAD) datasets, there is no risk for information leakage from the training to the test sets, which is a non-trivial problem [4]. Compared to homology-based unsupervised methods (such as EVE [4]), ESM1b and other protein language models have many compelling properties. Protein language models are simpler and faster to use since only a single AA sequence is required as input once a universal model has been trained (instead of training a separate model for each gene based on its distinct homologs). ESM1b also provides predictions for every possible variant, including those that reside within important protein domains not covered by MSA (**Fig. 1C-D**).

We leveraged our workflow to examine how alternative splicing influences variant effects. We highlighted 3,477 missense variants in ClinVar with high variance in predicted effects across isoforms (**Fig. 3**). Such variable predicted effects are common in many disease-causing genes, including *BRCA1, IRF6* and *TGFB3*. To study the phenomenon globally across the entire human genome, we denoted ∼1.8M variants in ∼9K genes as isoform sensitive (**Fig. 4A**). While isoform-sensitive variants tend to occur near splice sites and within genes containing domains disrupted by splicing (**Fig. 4B**), some splicing events are predicted to influence much larger or distant regions in the protein sequence (**Fig. 4C-D**).

We anticipate that our public resource would be useful for a broad range of human genetics tasks. For the diagnosis of Mendelian diseases, ESM1b effect scores could be integrated with other sources of information to resolve the ambiguity of VUS (**Fig. 2C-D**). Functionalization of VUS continues to serve a pressing need given their high prevalence in clinical sequencing efforts [10], which prevents many patients from receiving a clear diagnosis [2, 41–43]. For genetic association studies, effect scores could be used as priors for which variants are causal to improve the power of burden tests and resolution of statistical fine-mapping [1]. For protein engineering, it has been shown that ESM1b scores could be used to nominate gain-of-function variants that may confer therapeutic benefits [44]. Finally, using protein language models as variant functionalization tools could also inform basic research of protein function, including the identification of protein domains and other functional units (**Fig. 1D, Fig. 2G**) and discerning the functional differences between alternative isoforms (**Fig. 4**).

Over the past decades, computational variant functionalization tools have dramatically improved in accuracy and usability [4]. Given the results provided in this work, and in line with the track record of language models in protein research [18, 19, 25, 45] and machine learning in general [46], protein language models such as ESM1b currently seem to be one of the most promising approaches to determine the clinical and biological consequences of genetic variants. It has been shown that as language models scale in the number of parameters and training data, they tend to substantially improve [18, 46] (although this may not always be straightforward [47]). We expect that the same trend of larger, better protein language models will continue to benefit and improve variant effect prediction.

## Materials & Methods

### ESM1b

The code and pre-trained parameters for ESM1b [19] were taken from the model’s official GitHub repository at: https://github.com/facebookresearch/esm. Specifically, we used the esm1b_t33_650M_UR50S model.

### Handling long sequences

Because ESM1b uses learned positional embeddings (and self attention which grows quadratically with sequence length in both memory and compute), the model is restricted to protein sequences of up to 1,022aa in length [19]. However, ∼12% of human proteins in UniProt exceed this length [24]. To overcome this limitation, we used a sliding window approach where longer sequences were subdivided into overlapping windows of 1,022aa with at least 511aa overlap. More specifically, we tiled each full protein sequence with windows of size 1022aa starting from both ends. In every step, consecutive windows from both ends are generated to have an overlap of exactly 511aa. This process continues until the windows from both ends meet at the center. If the overlap between the central windows is less than 511aa, a last window of length 1022aa is added at the center to conclude the process. The subsequences defined by the windows are then used as inputs for ESM1b to get the LLR effect scores for all missense variants (each variant with respect to all the windows containing it). Since most residues are contained in multiple overlapping windows, the final effect score of each variant is determined by a weighted average approach. Specifically, to discount potential edge effects, the weights near window edges are constructed with a sigmoid function (**Supplementary Fig. S2**). The final variant effect score is then determined to be *(w(i1) * s1 + … + w(ik) * sk) / (w(i1) + … + w(ik))* where *s1,…,sk* are the effect scores of the variant in the context of each of the *k* windows containing it, *i1,…,ik* are the positions of the variant relative to each of these windows, and *w* is the window weight function (**Supplementary Fig. S2**).

### The global AUC metric for pathogenicity classification

To compare the performance of ESM1b and EVE [4] as variant pathogenicity classifiers, we used the ROC-AUC metric, which is the standard metric used to evaluate binary classifiers [48]. To calculate the ROC-AUC, we considered the entire set of pathogenic and benign variants (either from ClinVar [10] or HGMD/gnomAD [9, 26]) as a single genome-wide classification task, and consecutively calculated the global ROC-AUC of ESM1b and EVE over each of the two datasets (**Fig. 2B**). ESM1b is consistently superior to EVE according to this metric.

In the original publication introducing EVE [4], the authors also reported a somewhat different metric, which we refer to as gene-average AUC (as opposed to the global AUC reported in this work). Essentially, they treated each human gene as a separate dataset and evaluated EVE as a pathogenicity classifier over variants in that specific gene, calculating a gene-specific AUC for only 1,654 genes that have at least one annotated variant in each class (pathogenic/benign). They then averaged all the gene-specific AUC scores to obtain the final gene-average AUC. With respect to gene-average AUC on this smaller subset of genes, EVE is somewhat superior to ESM1b (**Supplementary Fig. S1**).

The fact that EVE is better on average across genes (gene-average AUC) but falls short of ESM1b at evaluating variant pathogenicity over all genes as a unified task (global AUC), suggests that ESM1b provides scores which are more consistent and comparable across different genes. This is somewhat expected given that EVE is in fact an assembly of multiple gene-specific models whereas ESM1b is a single model trained over the entire space of known protein sequences.

We argue that global AUC is a more appropriate metric than gene-average AUC for most purposes. From a clinical perspective, when attempting to diagnose a patient with a genetic disease, it is usually the case that variants across multiple genes are considered, and the overall evidence for each of the variants should be compared. In that case, it is important to obtain well-calibrated effect scores that can be compared between different genes, as evaluated by the global AUC metric.

### Funding

C.J.Y. is supported by the NIH grants R01AR071522, R01AI136972, U01HG012192, R01HG011239 and the Chan Zuckerberg Initiative, and is an investigator at the Chan Zuckerberg Biohub and is a member of the Parker Institute for Cancer Immunotherapy (PICI).

## Acknowledgements

We would like to thank Peter Stenson, Matthew Mort and David Cooper from Cardiff University for providing us with access to the HGMD database.

## Supplementary Figures & Tables

**Supplementary Figure S1.**
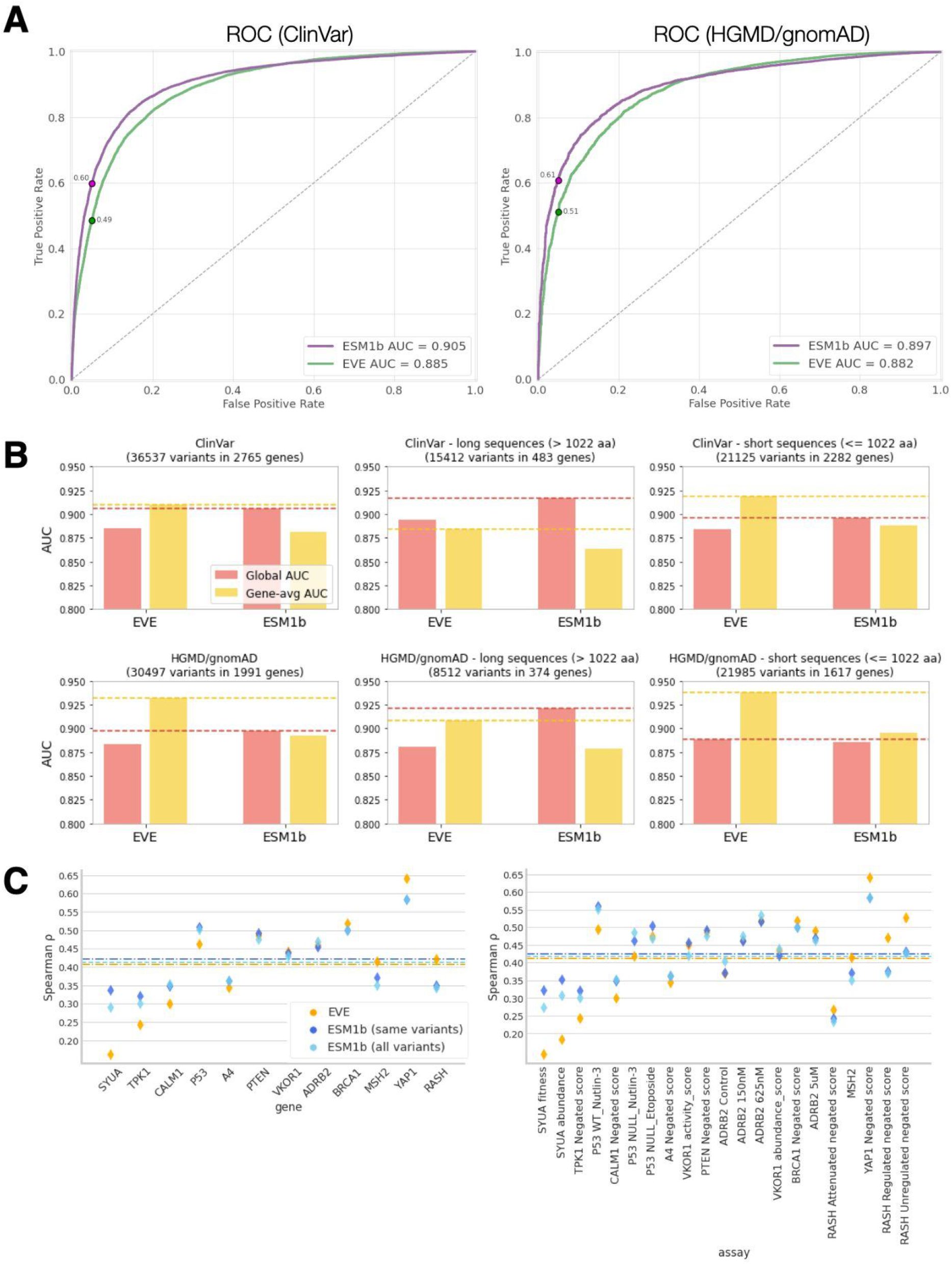
Comprehensive evaluation of ESM1b and EVE on ClinVar, HGMD/gnomAD and deep mutation scans. (**A**) ROC curves of ESM1b and EVE as binary classifiers of variant pathogenicity over ClinVar (left) and HGMD/gnomAD (right). The true positive rate at the standard false positive rate (0.05) is annotated across all 4 curves. (**B**) Evaluation of EVE (left bar plots) and ESM1b (right bar plots) over ClinVar (top panels) and HGMD/gnomAD (bottom panels), using either the global AUC (red) or gene-average AUC (yellow) metric (see the relevant section in the Methods). For each dataset, we show the results for either the full dataset (left panels), or the subsets of variants in long (middle panels) or short (right panels) proteins (defined by a threshold of 1,022aa, which is the maximum window length supported by ESM1b; see Methods). Dashed lines: the top score (obtained by ESM1b or EVE) according to each of the two metrics. (**C**) Evaluation of ESM1b and EVE on deep mutational scanning datasets. Right: Raw results over each of the 21 assays. Left: per-gene averages, across the 12 unique genes (as in Fig. 2E).

**Supplementary Figure S2.**
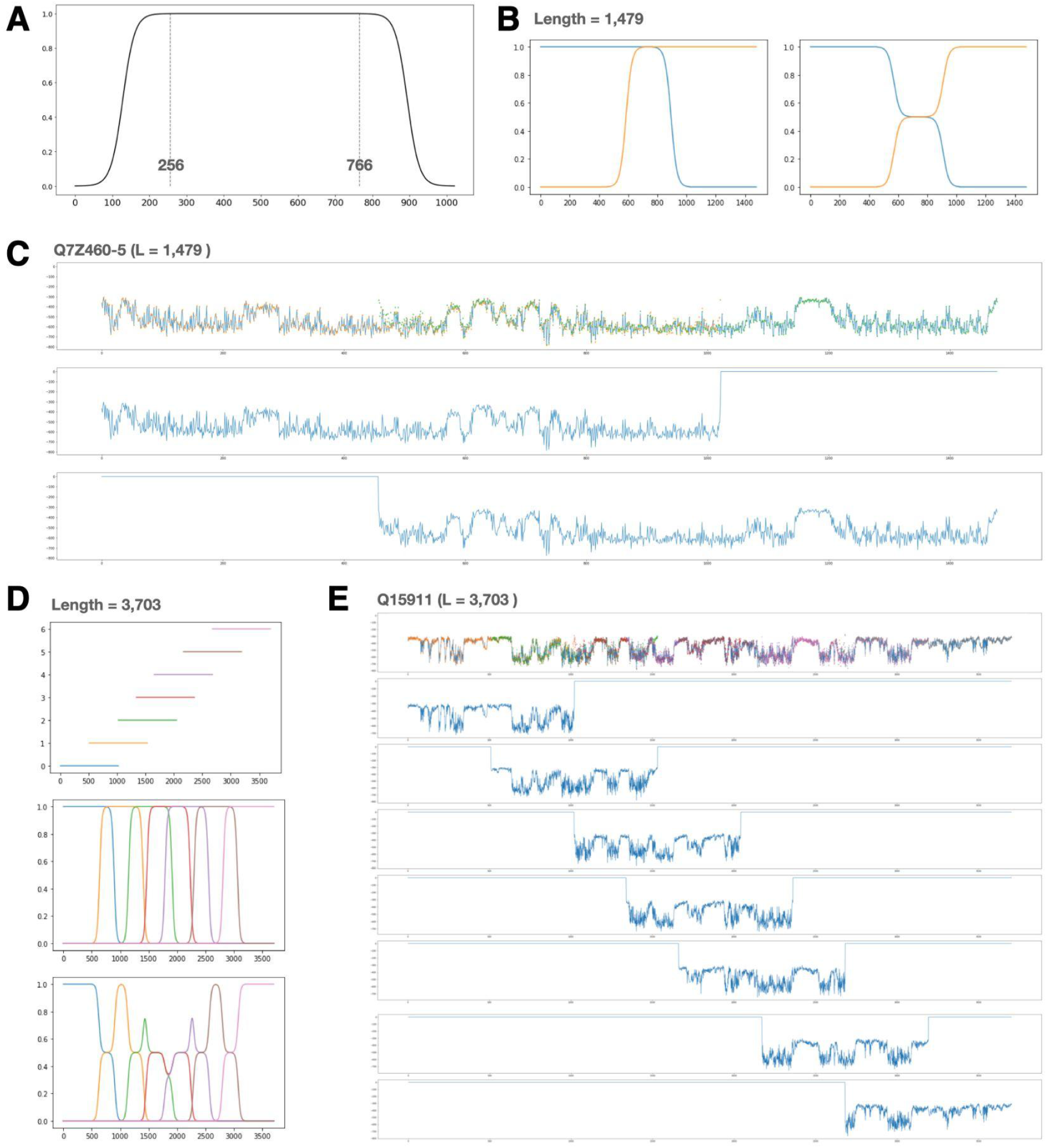
The sliding-window approach to tile long protein sequences with ESM1b. (**A**) The variant weights over each window’s coordinates (*1≤i≤1022*), defined by the function: *w(i) = 1 / (1 + exp(-(i-128)/16)* for *1≤i<256, w(i) = 1* for *256≤i<1022-256*, and *w(i) = 1/(1 + exp((i-1022+128)/16)* for *1022-256≤i≤1022*. (**B**) An example tiling of a protein sequence of length 1,479aa. Left: raw window weights (as in (A)). Right: normalized weights (summing up to 1 at each protein position). (**C**) Example of how a specific protein isoform (UniProt ID *Q7Z460-5*) is tiled. Top panel: ESM1b effect scores over the left window (*1≤i≤1022*; orange), the right window (*458≤i≤1479*; green), and the final weighted average throughout the entire protein’s length (blue). Middle: ESM1b effect scores over the left window. Bottom: ESM1b effect scores over the right window. (**D**) An example tiling of a larger protein sequence of length 3,703aa, as in (B). Top: the locations of the 7 windows used to tile the sequence. Middle: raw window weights. Bottom: normalized weights. (**E**) Example of how a specific protein (UniProt ID *Q15911*) is tiled, as in (C). As shown in the two examples, the effect scores tend to be consistent across different windows (with edge effects sometimes being more pronounced).

**Supplementary Table S1.**
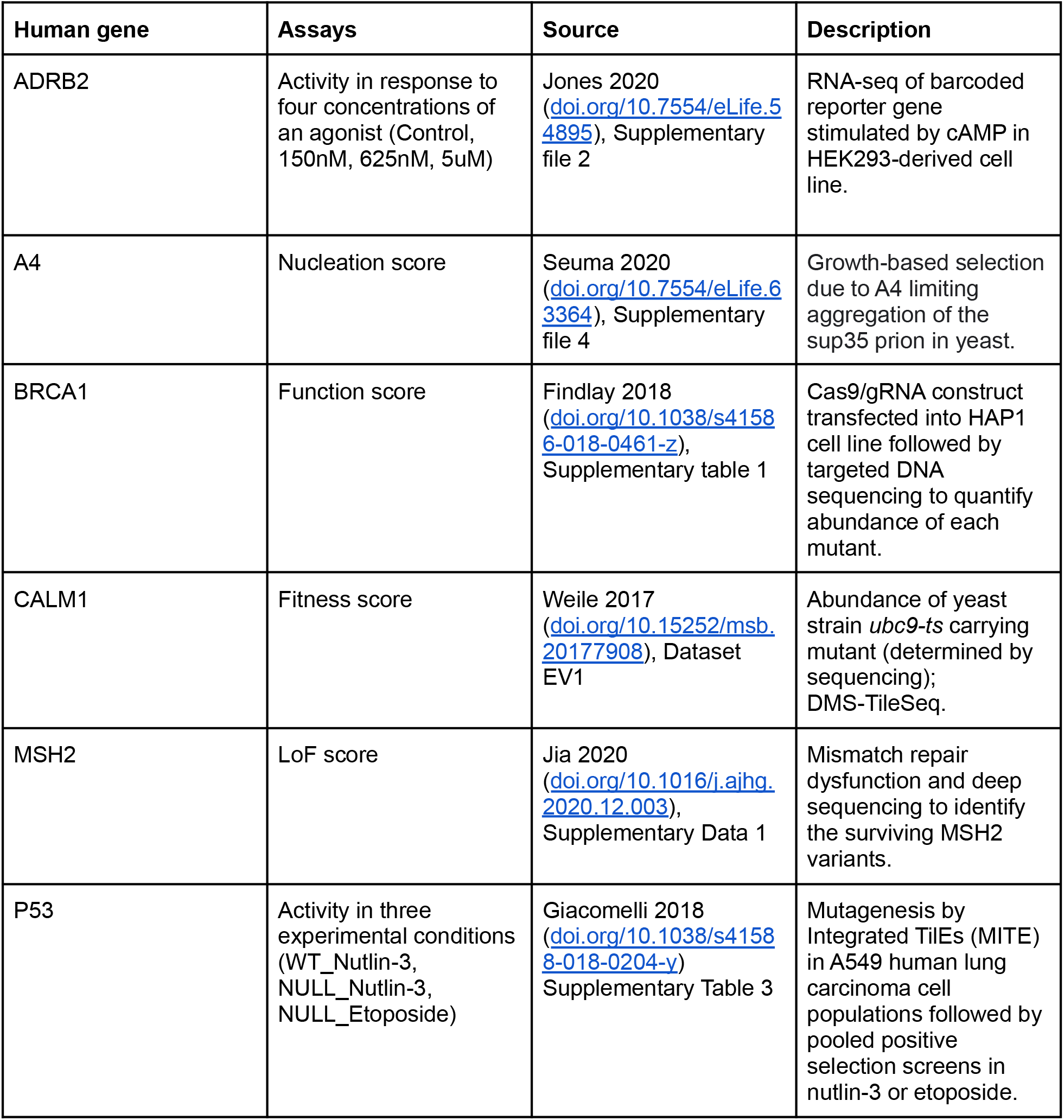
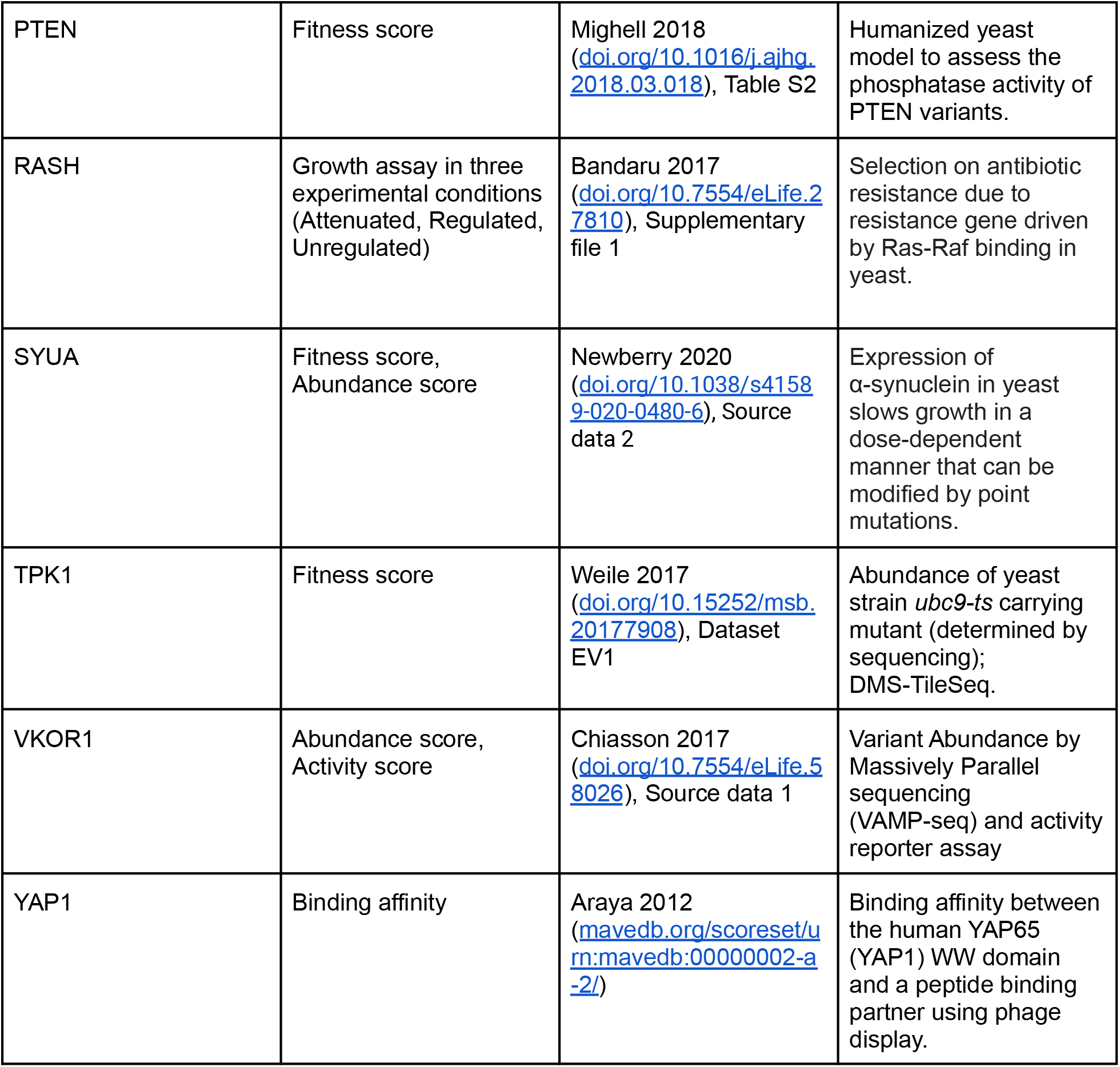
Deep mutational scans used for model evaluation.

## Supplementary Methods

### UniProt, ClinVar, HGMD and gnomAD datasets

The 42,336 protein isoform sequences were taken from UniProt [24] (in February 2022). To get ClinVar labels across all annotated variants and map them to UniProt protein sequences (**Fig. 2A, Fig. 2D** and **Fig. 3**), we downloaded the *variant_summary.txt.gz* file from ClinVar (on April 29, 2022). This dataset contained information about ∼1.3M variants, including clinical significance and NM code (RefSeq mRNA record). From a separate dataset on ClinVar’s website (*hgvs4variation.txt.gz*, downloaded on May 3, 2022), we obtained a mapping between the NM codes and NP codes (RefSeq protein records). We then matched those NP codes with UniProt IDs, using the same set of 42,336 protein isoforms from UniProt. Following this mapping, from NM code to NP code to UniProt ID, we obtained 977,833 variants with clinical significance contributing to 2,447,403 effects on UniProt isoforms (∼2.5 isoform effects per variant on average).

Access to the full HGMD dataset (https://www.hgmd.cf.ac.uk/ac/index.php) was provided upon request (see Acknowledgements) [26]. Of the entire set of 331,912 HGMD variants, 172,461 were missense variants, of which 158,725 were mapped to a unique UniProt isoform, of which 98,086 were annotated as disease-causing variants (DM).

To create a list of common missense variants, we used exomed-derived variants from the gnomAD database (version 2.1.1), by downloading the file *gnomad.exomes.r2.1.1.sites.vcf.bgz*. From this VCF file, we extracted 5,624,824 missense variants with a PASS filter, of which 4,876,367 were called in over 200,000 alleles (according to the AN value in the VCF file) and were considered high-quality variants. Of these, 26,359 variants (0.5%) had MAF > 0.01 and were considered common variants.

### Comparison to EVE, MSA coverage and deep mutational scans

EVE scores for missense variants affecting the ∼3K disease-associated genes analyzed by the model were downloaded from the official EVE portal (https://evemodel.org/). From these files we also extracted the ClinVar labels used for comparison between ESM1b and EVE (**Fig. 2B**), after filtering for high quality labels (at least 1 “Gold Stars”).

Multiple sequence alignments (MSAs) for ∼3K disease genes were downloaded from the EVE portal. For each MSA, the target (human) protein sequence is reported as a combination of uppercase and lowercase letters. The uppercase letters correspond to residues that are part of the MSA profile, whereas lowercase letters correspond to residues that are not. Accordingly, we defined the coverage of a human protein (reported in **Fig. 1C-D**) to be the fraction of the target sequence letters that are uppercase.

To compare the performance of ESM1b and EVE on deep mutational scanning experiments, we considered the same set of human genes as in [4] and downloaded the reported experimental data for all available assays (**Supplementary Table S1**). The only exception was Rhodopsin [49] for which the relevant data were not publicly available. For each assay we compared EVE and ESM1b on the same set of variants using the Spearman correlation between the predicted and experimental scores as the evaluation metric. Independent of EVE, we also evaluated performance of ESM1b on all experimental scores, including variants that were outside EVE/MSA coverage (**Fig. 2E** and **Supplementary Fig. S1**).

Throughout our evaluation, we used the raw experimental scores without any further processing for all deep mutational scans, except for CALM1, TPK1, RASH and the abundance assay of SYUA. For these assays, we transformed the scores using the following simple transformation: *x → |x - x_WT_|*, where *x*_*WT*_ is the assay-wide value measured for wild-type. The motivation for this transformation is that in most of these assays, variants that measure higher than the wild-type are considered deleterious for humans (see discussions in [50, 51]). In the case of SYUA, since variants that lead to decreased abundance are less toxic, their abundance scores were transformed the same way to better capture fitness (see Supplementary Figure 2 in [52]). Of note, this preprocessing significantly increased the evaluated performance of both EVE and ESM1b on these assays.

## References

1. Brandes N, Weissbrod O, Linial M (2022) Open problems in human trait genetics. Genome Biol 23:131. https://doi.org/10.1186/s13059-022-02697-9

2. Richards S, Aziz N, Bale S, et al (2015) Standards and guidelines for the interpretation of sequence variants: a joint consensus recommendation of the American College of Medical Genetics and Genomics and the Association for Molecular Pathology. Genet Med 17:405–423

3. Rehm HL, Fowler DM (2019) Keeping up with the genomes: scaling genomic variant interpretation. Genome Med 12:5. https://doi.org/10.1186/s13073-019-0700-4

4. Frazer J, Notin P, Dias M, et al (2021) Disease variant prediction with deep generative models of evolutionary data. Nature 1–5

5. Buniello A, MacArthur JAL, Cerezo M, et al (2018) The NHGRI-EBI GWAS Catalog of published genome-wide association studies, targeted arrays and summary statistics 2019. Nucleic Acids Res 47:D1005—-D1012

6. Hamosh A, Scott AF, Amberger JS, et al (2005) Online Mendelian Inheritance in Man (OMIM), a knowledgebase of human genes and genetic disorders. Nucleic Acids Res 33:D514—-D517

7. Finucane HK, Bulik-Sullivan B, Gusev A, et al (2015) Partitioning heritability by functional annotation using genome-wide association summary statistics. Nat Genet 47:1228–1235. https://doi.org/10.1038/ng.3404

8. Brandes N, Linial N, Linial M (2021) Genetic association studies of alterations in protein function expose recessive effects on cancer predisposition. Sci Rep 11:14901. https://doi.org/10.1038/s41598-021-94252-y

9. Gudmundsson S, Singer-Berk M, Watts NA, et al (2021) Variant interpretation using population databases: Lessons from gnomAD. Hum Mutat

10. Landrum MJ, Lee JM, Benson M, et al (2015) ClinVar: public archive of interpretations of clinically relevant variants. Nucleic Acids Res 44:D862—-D868

11. Esposito D, Weile J, Shendure J, et al (2019) MaveDB: an open-source platform to distribute and interpret data from multiplexed assays of variant effect. Genome Biol 20:223. https://doi.org/10.1186/s13059-019-1845-6

12. Ursu O, Neal JT, Shea E, et al (2022) Massively parallel phenotyping of coding variants in cancer with Perturb-seq. Nat Biotechnol. https://doi.org/10.1038/s41587-021-01160-7

13. Boucher JI, Bolon DN, Tawfik DS (2016) Quantifying and understanding the fitness effects of protein mutations: Laboratory versus nature. Protein Sci 25:1219–1226

14. Hopf TA, Ingraham JB, Poelwijk FJ, et al (2017) Mutation effects predicted from sequence co-variation. Nat Biotechnol 35:128–135

15. Ng PC (2003) SIFT: predicting amino acid changes that affect protein function. Nucleic Acids Res 31:3812–3814. https://doi.org/10.1093/nar/gkg509

16. Adzhubei I, Jordan DM, Sunyaev SR (2013) Predicting functional effect of human missense mutations using PolyPhen-2. Curr Protoc Hum Genet 76:7–20

17. Rentzsch P, Witten D, Cooper GM, et al (2019) CADD: predicting the deleteriousness of variants throughout the human genome. Nucleic Acids Res 47:D886–D894

18. Ofer D, Brandes N, Linial M (2021) The language of proteins: NLP, machine learning & protein sequences. Comput Struct Biotechnol J

19. Rives A, Meier J, Sercu T, et al (2021) Biological structure and function emerge from scaling unsupervised learning to 250 million protein sequences. Proc Natl Acad Sci 118:

20. Elnaggar A, Ding W, Jones L, et al (2021) CodeTrans: Towards Cracking the Language of Silicon’s Code Through Self-Supervised Deep Learning and High Performance Computing. ArXiv Prepr ArXiv210402443

21. Strodthoff N, Wagner P, Wenzel M, Samek W (2020) UDSMProt: universal deep sequence models for protein classification. Bioinformatics 36:2401–2409

22. Alley EC, Khimulya G, Biswas S, et al (2019) Unified rational protein engineering with sequence-based deep representation learning. Nat Methods 16:1315–1322

23. Brandes N, Ofer D, Peleg Y, et al (2022) ProteinBERT: a universal deep-learning model of protein sequence and function. Bioinformatics 38:2102–2110. https://doi.org/10.1093/bioinformatics/btac020

24. Boutet E, Lieberherr D, Tognolli M, et al (2016) UniProtKB/Swiss-Prot, the manually annotated section of the UniProt KnowledgeBase: how to use the entry view. In: Plant Bioinformatics. Springer, pp 23–54

25. Meier J, Rao R, Verkuil R, et al (2021) Language models enable zero-shot prediction of the effects of mutations on protein function. bioRxiv

26. Stenson PD, Ball EV, Mort M, et al (2003) Human gene mutation database (HGMD®): 2003 update. Hum Mutat 21:577–581

27. Allison R, Edgar JR, Reid E (2019) Spastin MIT domain disease-associated mutations disrupt lysosomal function. Front Neurosci 13:1179

28. Nava C, Rupp J, Boissel J-P, et al (2015) Hypomorphic variants of cationic amino acid transporter 3 in males with autism spectrum disorders. Amino Acids 47:2647–2658

29. Shoubridge C, Tan MH, Seiboth G, Gecz J (2012) ARX homeodomain mutations abolish DNA binding and lead to a loss of transcriptional repression. Hum Mol Genet 21:1639–1647

30. Bienvenu T, Poirier K, Friocourt G, et al (2002) ARX, a novel Prd-class-homeobox gene highly expressed in the telencephalon, is mutated in X-linked mental retardation. Hum Mol Genet 11:981–991

31. Marques I, Sá MJ, Soares G, et al (2015) Unraveling the pathogenesis of ARX polyalanine tract variants using a clinical and molecular interfacing approach. Mol Genet Genomic Med 3:203–214

32. Cho G, Nasrallah MP, Lim Y, Golden JA (2012) Distinct DNA binding and transcriptional repression characteristics related to different ARX mutations. neurogenetics 13:23–29

33. Huang J, Gurung B, Wan B, et al (2012) The same pocket in menin binds both MLL and JUND but has opposite effects on transcription. Nature 482:542–546

34. Cebrian A, Ruiz-Llorente S, Cascon A, et al (2003) Mutational and gross deletion study of the MEN1 gene and correlation with clinical features in Spanish patients. J Med Genet 40:e72–e72

35. Martín-Campos JM, Catasús L, Chico A, et al (1999) Molecular pathology of multiple endocrine neoplasia type I: two novel germline mutations and updated classification of mutations affecting MEN1 gene. Diagn Mol Pathol Am J Surg Pathol Part B 8:195–204

36. Agarwal SK, Guru SC, Heppner C, et al (1999) Menin interacts with the AP1 transcription factor JunD and represses JunD-activated transcription. Cell 96:143–152

37. Klein RD, Salih S, Bessoni J, Bale AE (2005) Clinical testing for multiple endocrine neoplasia type 1 in a DNA diagnostic laboratory. Genet Med 7:131–138

38. Toledo RA, Lourenco DM, Coutinho FL, et al (2007) Novel MEN1 germline mutations in Brazilian families with multiple endocrine neoplasia type 1. Clin Endocrinol (Oxf) 67:377–384

39. Ehrlich L, Hall C, Venter J, et al (2017) miR-24 inhibition increases menin expression and decreases cholangiocarcinoma proliferation. Am J Pathol 187:570–580

40. Jumper J, Evans R, Pritzel A, et al (2021) Highly accurate protein structure prediction with AlphaFold. Nature 596:583–589

41. Nicora G, Zucca S, Limongelli I, et al (2022) A machine learning approach based on ACMG/AMP guidelines for genomic variant classification and prioritization. Sci Rep 12:2517. https://doi.org/10.1038/s41598-022-06547-3

42. Tavtigian SV, Greenblatt MS, Harrison SM, et al (2018) Modeling the ACMG/AMP variant classification guidelines as a Bayesian classification framework. Genet Med 20:1054–1060

43. Tavtigian SV, Harrison SM, Boucher KM, Biesecker LG (2020) Fitting a naturally scaled point system to the ACMG/AMP variant classification guidelines. Hum Mutat 41:1734–1737

44. Hie BL, Xu D, Shanker VR, et al (2022) Efficient evolution of human antibodies from general protein language models and sequence information alone. bioRxiv

45. Rao R, Bhattacharya N, Thomas N, et al (2019) Evaluating protein transfer learning with tape. Adv Neural Inf Process Syst 32:9689

46. Thoppilan R, De Freitas D, Hall J, et al (2022) Lamda: Language models for dialog applications. ArXiv Prepr ArXiv220108239

47. Nijkamp E, Ruffolo J, Weinstein EN, et al (2022) ProGen2: Exploring the Boundaries of Protein Language Models. ArXiv Prepr ArXiv220613517

48. Pedregosa F, Varoquaux G, Gramfort A, et al (2011) Scikit-learn: Machine learning in Python. J Mach Learn Res 12:2825–2830

49. Penn WD, McKee AG, Kuntz CP, et al (2020) Probing biophysical sequence constraints within the transmembrane domains of rhodopsin by deep mutational scanning. Sci Adv 6:eaay7505

50. Weile J, Sun S, Cote AG, et al (2017) A framework for exhaustively mapping functional missense variants. Mol Syst Biol 13:957

51. Bandaru P, Shah NH, Bhattacharyya M, et al (2017) Deconstruction of the Ras switching cycle through saturation mutagenesis. Elife 6:e27810

52. Newberry RW, Leong JT, Chow ED, et al (2020) Deep mutational scanning reveals the structural basis for α-synuclein activity. Nat Chem Biol 16:653–659

